# Broad-spectrum in vitro antiviral activity of ODBG-P-RVn: an orally-available, lipid-modified monophosphate prodrug of remdesivir parent nucleoside (GS-441524)

**DOI:** 10.1101/2021.08.06.455494

**Authors:** Michael K. Lo, Punya Shrivastava-Ranjan, Payel Chatterjee, Mike Flint, James R. Beadle, Nadejda Valiaeva, Robert T. Schooley, Karl Y. Hostetler, Joel M. Montgomery, Christina Spiropoulou

## Abstract

The intravenous administration of remdesivir for COVID-19 confines its utility to hospitalized patients. We evaluated the broad-spectrum antiviral activity of ODBG-P-RVn, an orally available, lipid-modified monophosphate prodrug of the remdesivir parent nucleoside (GS-441524) against viruses that cause diseases of human public health concern, including SARS-CoV-2. ODBG-P-RVn showed 20-fold greater antiviral activity than GS-441524 and had near-equivalent activity to remdesivir in primary-like human small airway epithelial cells. Our results warrant investigation of ODBG-P-RVn efficacy in vivo.

---

Remdesivir (RDV; Veklury, GS-5734) is an adenosine nucleotide analog phosphoramidate prodrug with broad-spectrum antiviral activity in vitro and in vivo (1–8), and is currently the only therapeutic approved by the FDA for treating coronavirus 19 disease (COVID-19) in hospitalized patients over the age of 12 (9). While RDV did not significantly reduce COVID-19 mortality, it did shorten the time to recovery compared to a placebo control group (10). The short half-life of RDV in human and animal plasma (1, 8, 11, 12), alongside the in vivo efficacy of RDV parent nucleoside (GS-441524, RVn) against coronaviruses including severe acute respiratory syndrome coronavirus 2 (SARS-CoV-2) (13–16), have driven proposals to utilize RVn instead of RDV to treat COVID-19 (17). A recent comparative pharmacokinetic study in non-human primates, however, demonstrated higher nucleoside triphosphate (NTP) levels in lower respiratory tract tissues of RDV-dosed animals than in RVn-dosed animals (8). A significant drawback of RDV is the requirement for intravenous administration, which limits its use to hospital contexts. In an attempt to develop an orally bioavailable form of remdesivir, we recently synthesized a 1-O-octadecyl-2-O-benzyl-sn-glycerylester (ODBG) lipid-modified monophosphate prodrug of RVn (ODBG-P-RVn), which demonstrated more favorable in vitro antiviral activity against SARS-CoV-2 compared to that of RVn and RDV in Vero-E6 cells (18).

In this study, we extended our in vitro comparisons to include 14 viruses from across 7 virus families responsible for causing diseases of significant human public health concern. These were *Filoviridae*: Ebola virus (EBOV) and Marburg virus (MARV) (19, 20); *Paramyxoviridae*: Nipah virus (NiV), Hendra virus (HeV), human parainfluenza virus 3 (hPIV3), measles virus (MV), mumps virus (MuV), and Sosuga virus (SoSuV) (21–27); *Pneumoviridae*: respiratory syncytial virus (RSV) (28); *Flaviviridae*: yellow fever virus (YFV); *Arenaviridae*: Lassa virus (LASV) (29); *Nairoviridae*: Crimean-Congo hemorrhagic fever virus (CCHFV) (30); and *Coronaviridae*: SARS-CoV-2 (31). We utilized 3 previously described assays to compare the antiviral activities of RVn, RDV, and ODBG-P-RVn against this panel of viruses: 1) directly measuring fluorescence of a reporter protein expressed by recombinant viruses (REP) (2), (Figure 1A); 2) quantitating focus-forming units (FFU) via fluorescent reporter imaging (32) (Figure 1B); and 3) indirectly measuring cytopathic effect (CPE) based on cellular ATP levels (CellTiterGlo 2.0, Promega) (2) (Figure 1C), which was also used to evaluate compound cytotoxicity (Figure 1D). Assay conditions varied based on virus replication kinetics and on the specific assay used; multiplicities of infection (MOI) ranged from 0.01–0.25, and endpoint measurements were conducted between 72-144 hours post-infection (hpi). We initially conducted dose-response experiments using 8-point, 3-fold serial dilutions of RVn, RDV, and ODBG-P-RVn against our panel of viruses in Vero-E6 cells, and showed that ODBG-P-RVn consistently had greater antiviral activity than RVn and RDV against all viruses susceptible to RVn/RDV inhibition, with effective concentration (EC_50_) values ranging from 0.026 to 1.13 μM (Figure 1, Vero-E6 assays represented in left column of panels A, B, C; Supplemental Figure S1; Table 1). RVn and ODBG-P-RVn induced partial cytotoxicity but only at the highest concentration tested (100 μM) and without reaching 50% cytotoxicity (CC_50_). We then compared these antivirals in human hepatoma (Huh7) and bronchioalveolar carcinoma (NCI-H358) cell lines, which represent more relevant cell types targeted by subsets of viruses used in our study. In both human cell lines, although ODBG-P-RVn showed EC_50_ values remarkably similar to those observed in Vero-E6 cells and was 3- to 5-fold more active than RVn, it consistently showed 6- to 20-fold less activity than RDV (Figure 1 [Huh7 and NCI-H358 assays represented, respectively, in the middle and right columns of panels A, B, and C]; Supplemental Figures S2, S3; Table 1). Whereas CC_50_ values for RDV in Huh7 and NCI-H358 cells were 54.2 and 77.2 μM, respectively, ODBG-P-RVn was less cytotoxic in Huh7 cells (CC_50_ = 93.4 μM) and did not show measurable cytotoxicity in NCI-H358 cells even at the highest concentration tested (100 μM) (Figure 1D, right panel; Table 1).

**Figure 1.**
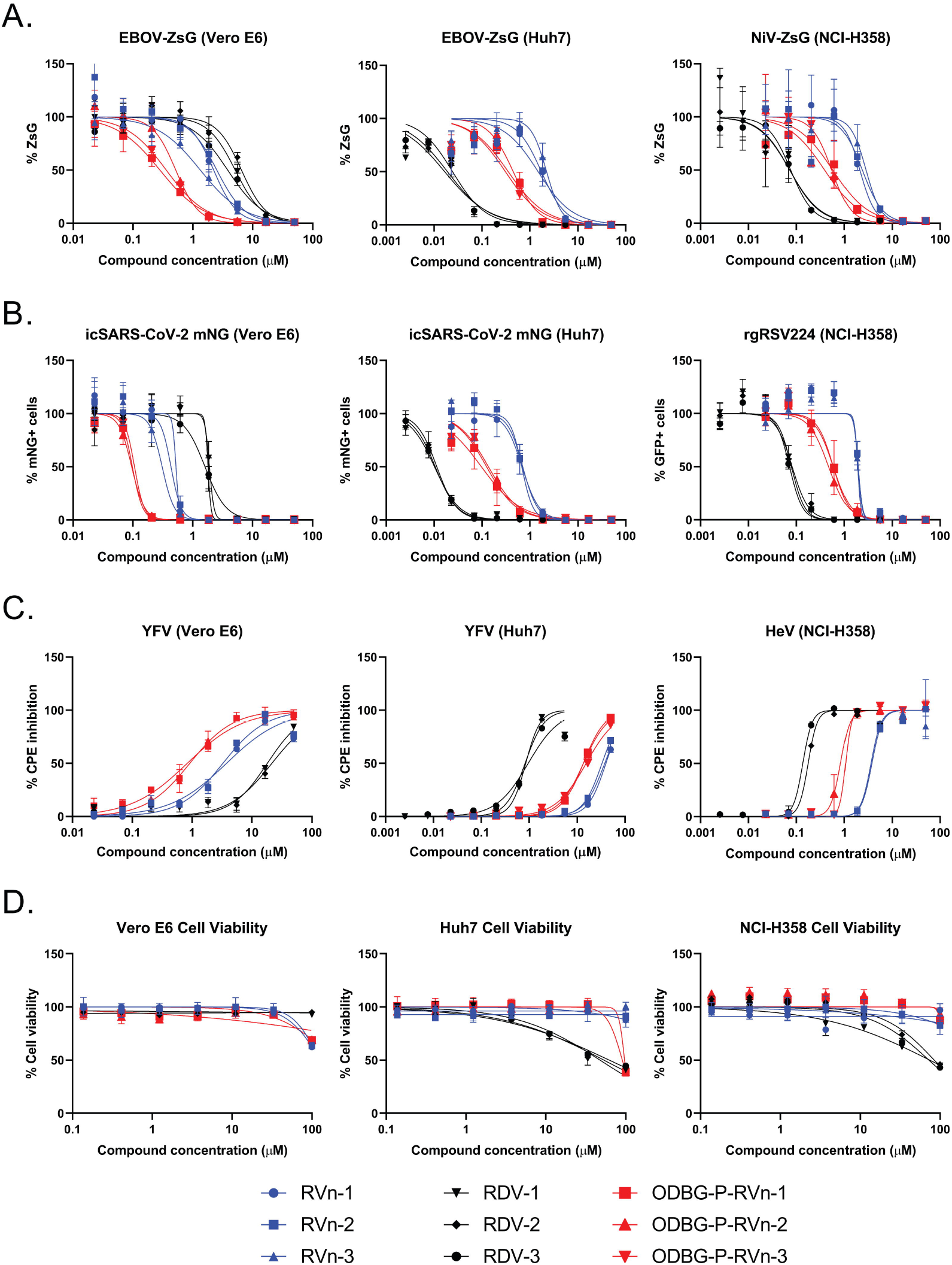
Comparison of antiviral activities of RVn, RDV, and ODBG-P-RVn in African green monkey (Vero-E6), human hepatoma (Huh7), and human bronchioalveolar carcinoma (NCI-H358) cell lines using reporter-based, image-based, and cytopathic effect (CPE) assays. Representative dose-response inhibition of viral replication and induction of cellular cytotoxicity by RVn (blue shapes), RDV (black shapes), and ODBG-P-RVn (red shapes). A) Direct measurement of reporter fluorescence intensity by recombinant Ebola virus (EBOV) expressing ZsGreen protein in Vero-E6 (left panel) and Huh7 (middle panel) cells, and recombinant Nipah virus (NiV) expressing ZsGreen protein in NCI-H358 (right panel) cells. B) Image-based counting of reporter fluorescence-positive cells infected with recombinant severe acute respiratory syndrome coronavirus 2 (SARS-CoV-2) expressing mNeonGreen protein (Vero-E6 and Huh7) and recombinant respiratory syncytial virus (RSV) expressing eGFP (NCI-H358). Infected cells treated with DMSO were considered as 100% fluorescence intensity signal and 100% fluorescence-positive cell counts. C) Compound-based inhibition of CPE induced by yellow fever virus (YFV) in Vero-E6 and Huh7 cells and by Hendra virus (HeV) in NCI-H358 cells determined by measuring cellular ATP levels (CellTiterGlo 2.0). ATP levels in uninfected cells treated with DMSO were considered 100% CPE inhibition. D) Compound cytotoxicity/cell viability measured by CellTiterGlo 2.0 assay. Dose-response curves were fitted to the mean value of experiments performed in biological triplicate for each concentration in the 8-point, 3-fold dilution series using a 4-parameter non-linear logistic regression curve with variable slope. Data points and error bars indicate the mean value and standard deviation of 3 biological replicates; each colored shape/line in the legend represents an independent experiment performed in biological triplicate. RVn and RDV used in this study was obtained from MedChemExpress (Monmouth Junction, NJ USA).

**Table 1.**
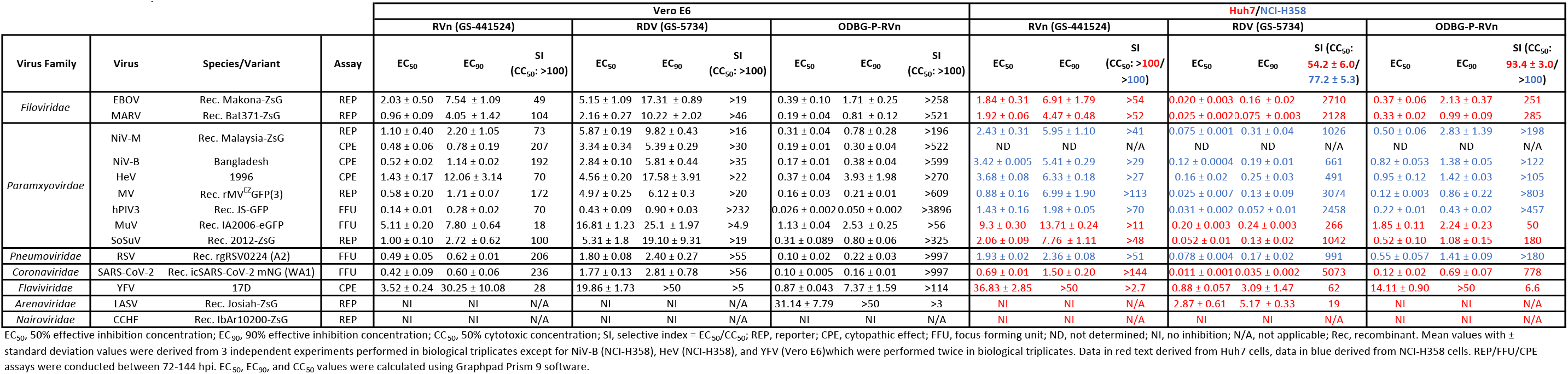
Mean antiviral activity of RVn, RDV, and ODBG-P-RVn in Vero E6, Huh7, and NCI-H358 cell lines.

To further evaluate cell type-specific effects on the antiviral activities of RVn, RDV, and ODBG-P-RVn, we tested them against a smaller subset of filoviruses (EBOV-ZsG, MARV-ZsG) and a paramyxovirus (NiV-ZsG) expressing ZsGreen reporter in primary-like human telomerase reverse transcriptase (hTERT) immortalized human microvascular endothelial (TIME) cells (33, 34). In TIME cells, we observed a similar trend in antiviral activity as in Huh7 and NCI-H358 cells, with ODBG-P-RVn showing 15- to 22-fold greater activity than RVn, but 5- to 8-fold less activity than RDV in reporter-based assays (Figure 2A, Table 2). To confirm this, we compared the respective abilities of RDV and ODBG-P-RVn to reduce infectious yield of EBOV-ZsG and NiV-ZsG (MOI = 0.25) when cells were treated with each compound 2 hpi. Virus supernatants were collected at 72hpi and titered on Huh7 (for EBOV-ZsG) or NCI-H358 (for NiV-ZsG) cells to determine 50% tissue culture infectious dose (TCID_50_) by the method of Reed and Muench (35). Both RDV and ODBG-P-RVn equivalently reduced infectious yield of EBOV-ZsG by up to 4 log_10_ and of NiV-ZsG by approximately 2 log_10_, in a dose-dependent manner, with EC_50_ values closely mirroring values determined in reporter assays (Figure 2B, left and middle panels; Table 2). However, RDV was more cytotoxic (CC_50_ = 17.2 μM) than ODBG-P-RVn (CC_50_ > 50 μM) (Figure 2B, right panel; Table 2), which is reflected in its biphasic inhibition of NiV-ZsG (Figure 2B, middle panel, cytotoxic inhibition by RDV shown at 16.6 μM). Since the ODBG lipid modification has been shown to enhance in vivo lung tissue distribution for a different orally administered nucleoside (36), we compared the activity of the 3 compounds against filoviruses, paramyxoviruses, and RSV in another primary-like, hTERT-immortalized small airway epithelial cell (HSAEC1-KT) (37). Notably, the dose-response curves of RDV and ODBG-P-RVn were strikingly similar, with EC_50_ values in the submicromolar range within a 3-fold range of each other; EC_50_ values for some viruses were almost identical (Figure 2C; Supplemental Figure 4; Table 2). Furthermore, RDV and ODBG-P-RVn equivalently reduced the infectious yields of EBOV-ZsG and NiV-ZsG in HSAEC1-KT cells by by 5 log_10_ and 3 log_10_, respectively, and their EC_50_ values reflected the limited differential in antiviral activity between them (Figure 2D, left and middle panels; Table 2). Although ODBG-P-RVn was more cytotoxic (CC_50_ = 20.5) in HSAEC1-KT cells than RDV (CC_50_ > 100; Figure 2D, right panel; Table 2), it also effectively reduced virus yields at non-cytotoxic concentrations.

**Figure 2.**
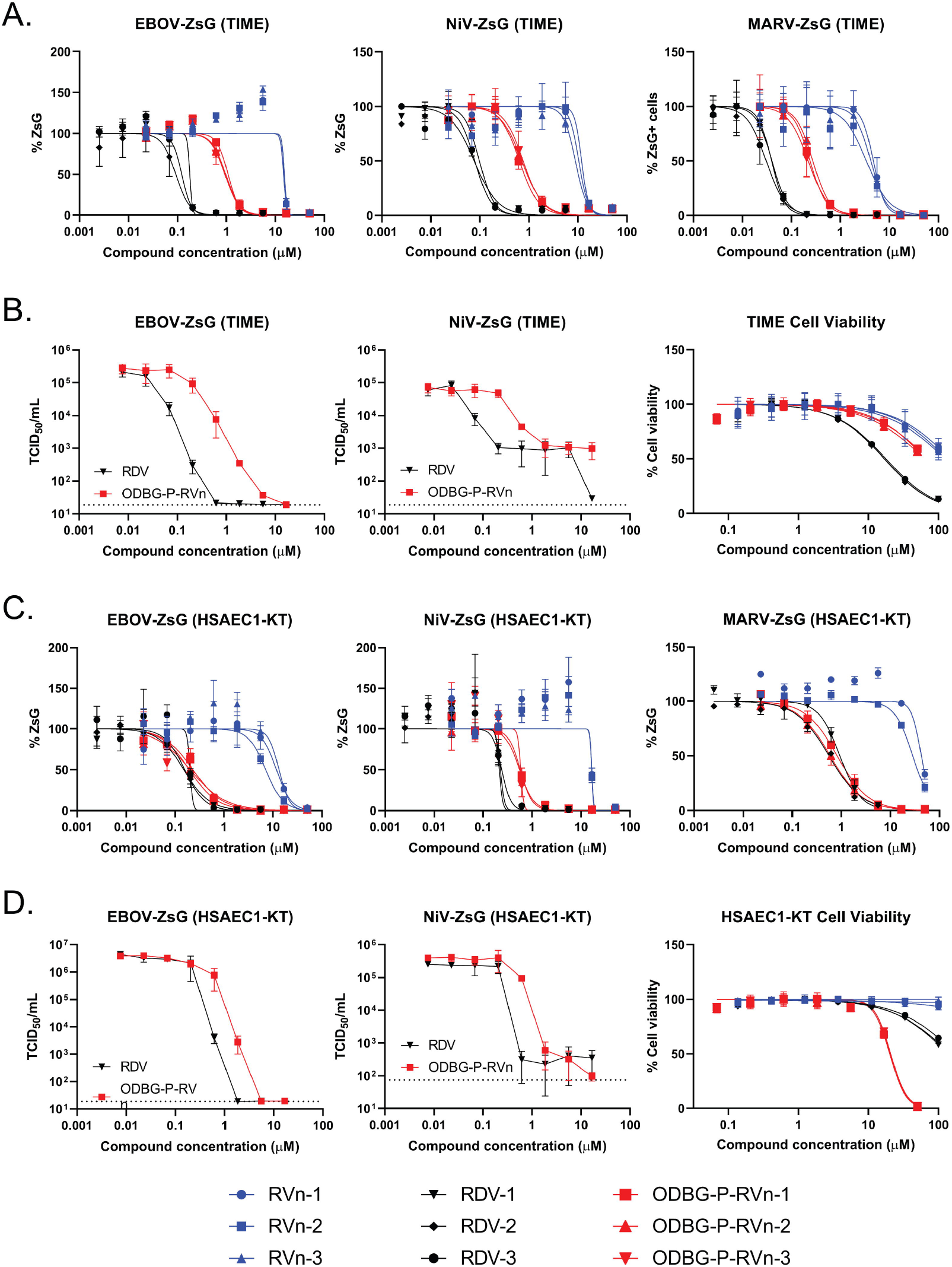
Comparison of cell type-dependent antiviral activities of RVn, RDV, and ODBG-P-RVn in primary-like hTERT-immortalized microvascular endothelial (TIME) cells and small airway epithelial cells (HSAEC1-KT). A) Representative dose-response inhibition of recombinant EBOV, NiV, and Marburg virus (MARV) expressing ZsGreen protein in TIME cells. B) Yield reduction of infectious EBOV-ZsG (left panel) and NiV-ZsG (middle panel) by RDV and ODBG-P-RVn. Compound cytotoxicity/cell viability (right panel) in TIME cells measured via CellTiterGlo 2.0 assay. C) Representative dose-response inhibition of recombinant EBOV, NiV, and MARV expressing ZsGreen protein in HSAEC1-KT cells. D) Reduction of infectious yield of EBOV-ZsG (left panel) and NiV-ZsG (middle panel) by RDV and ODBG-P-RVn in HSAEC1-KT cells. Compound cytotoxicity/cell viability (right panel) in HSAEC1-KT cells measured via CellTiterGlo 2.0 assay. Dose-response curves were fitted to the mean value of experiments performed in biological triplicate for each concentration in the 8-point, 3-fold dilution series using a 4-parameter non-linear logistic regression curve with variable slope. Data points and error bars indicate the mean value and standard deviation of 3 or 4 biological replicates; each colored shape/line in the legend represents an independent experiment performed in biological triplicate. Infectious yield reduction assays were conducted once with biological quadruplicates.

**Table 2.**
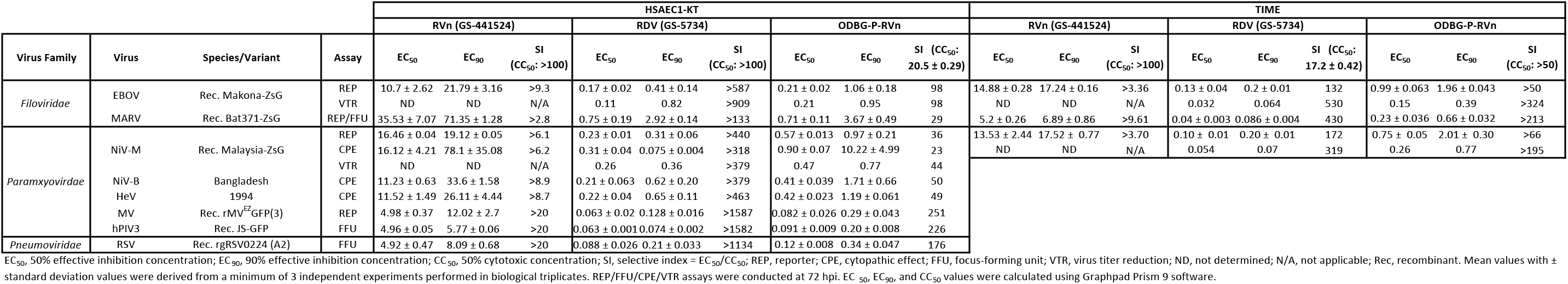
Mean antiviral activity of RVn, RDV, and ODBG-P-RVn in primary-like hTERT-immortalized microvascular endothelial (TIME) and small airway epithelial (HSAEC1-KT) cell lines.

In summary, our results demonstrate that ODBG-P-RVn has greater antiviral activity than RVn in all cell lines tested and has cell-type dependent activity levels that range from moderately lesser than to nearly equal to those of RDV. *In vivo* RDV is converted rapidly to RVn (1, 8, 11, 12), which has 0.5 to 2 log_10_ less activity than RDV against most of the viruses tested. In contrast, ODBG-P-RVn is stable in plasma for >24 hours and at therapeutic plasma levels of ODBG-P-Rvn (above EC_90_ for SARS-CoV-2) after oral administration of 16.9 mg/kg to Syrian hamsters; furthermore RVn was not observed at virologically significant levels (38). Thus, one would predict sustained *in vivo* antiviral activity with ODBG-P-RVn without substantial generation in plasma of RVn, the less active metabolite. Taken together, our results strongly support investigation of in vivo efficacy of ODBG-P-RVn not only against SARS-CoV-2 but also against other viruses significant to human health.

## ACKNOWLEDGMENTS

We thank Tatyana Klimova for helpful comments in reviewing the manuscript. We thank Pei-Yong Shi (University of Texas Medical Branch) for the kind gift of the reporter SARS-CoV-2 expressing mNeonGreen. The findings and conclusions in this report are those of the authors and do not necessarily represent those of the Centers for Disease Control and Prevention. This work was supported by CDC core funding and by the National Institute of Allergy and Infectious Diseases (RO1-AI131424).

## SUPPLEMENTAL FIGURE LEGENDS

**Supplemental Figure S1.**
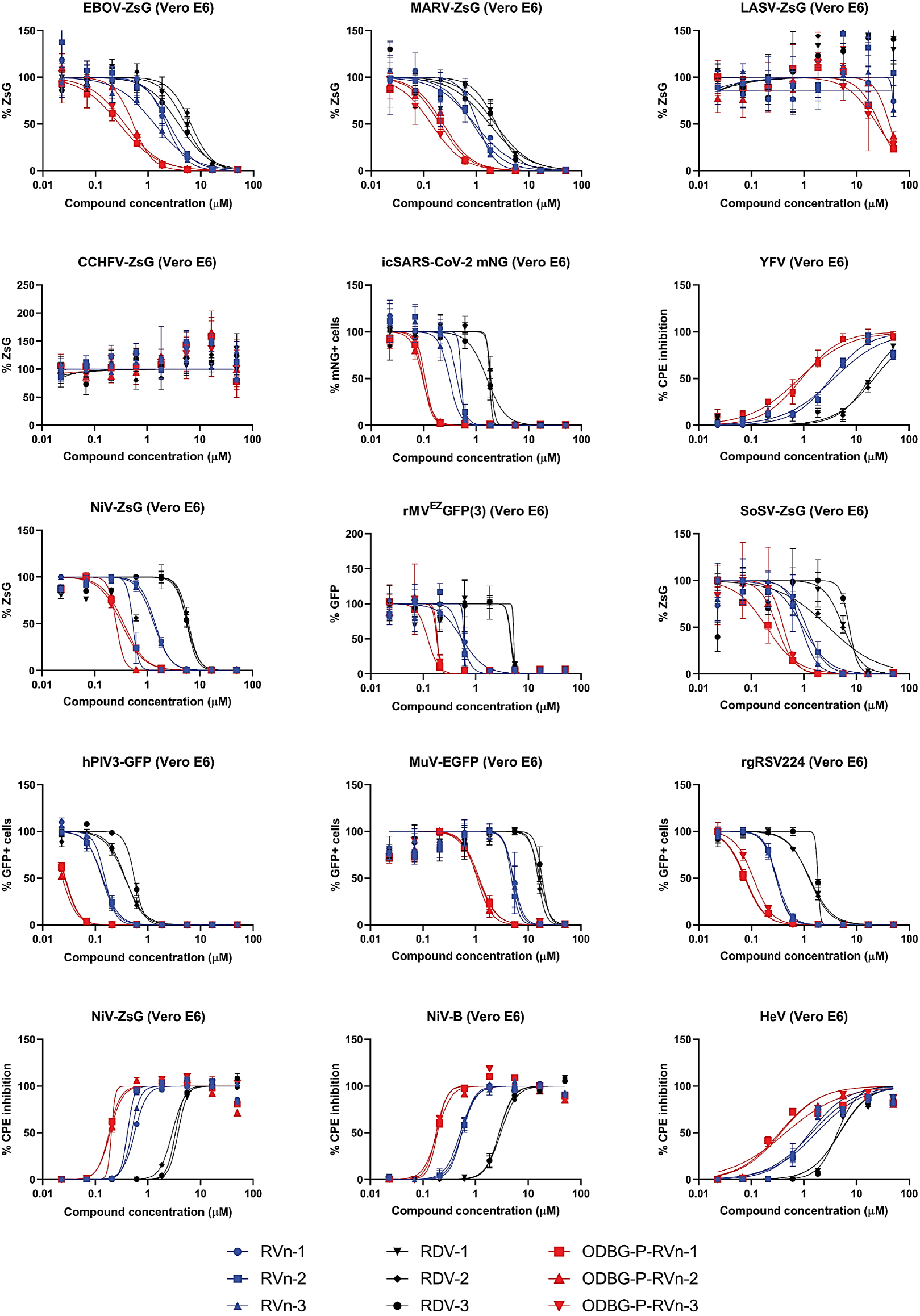
Comparison of antiviral activities of RVn, RDV, and ODBG-P-RVn in African green monkey (Vero-E6) cells using reporter-based, image-based, and CPE assays. Representative dose-response inhibition of virus replication by RVn (blue shapes), RDV (black shapes), and ODBG-P-RVn (red shapes). Signal from infected cells treated with DMSO served as 100% fluorescence intensity signal for reporter assays and 100% fluorescence-positive cell counts for image-based assays. CPE inhibition was measured by determining cellular ATP levels using CellTiterGlo 2.0 assay reagent. ATP levels in uninfected cells treated with DMSO served as 100% CPE inhibition. Dose-response curves were fitted to the mean value of experiments performed in biological triplicate for each concentration in the 8-point, 3-fold dilution series using a 4-parameter non-linear logistic regression curve with variable slope. Data points and error bars indicate the mean value and standard deviation of 3 biological replicates; each colored shape/line in the legend represents an independent experiment performed in biological triplicate.

**Supplemental Figure S2.**
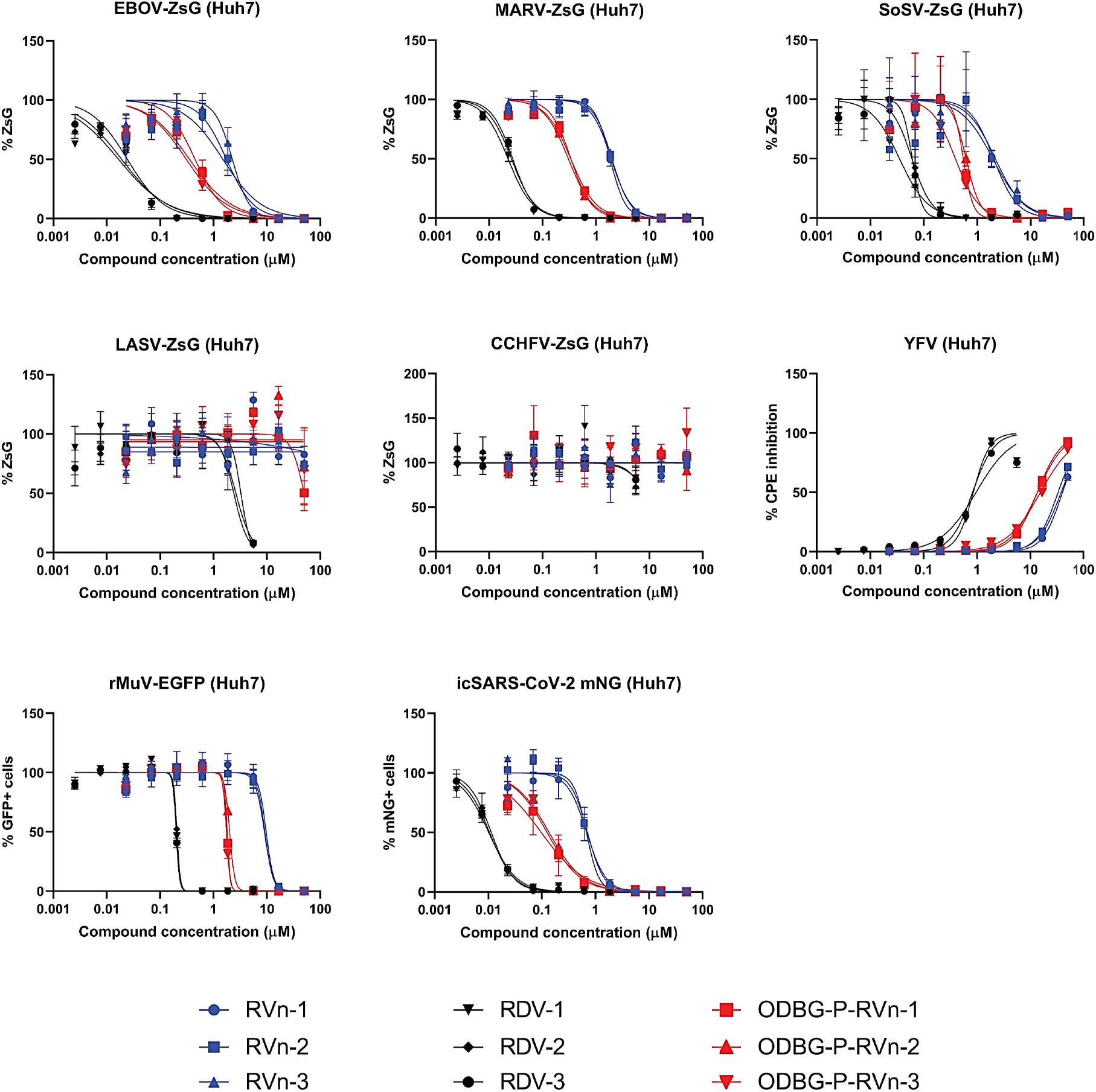
Comparison of antiviral activities of RVn, RDV, and ODBG-P-RVn in Huh7 cells using reporter-based, image-based, and CPE assays. Representative dose-response inhibition of virus replication by RVn (blue shapes), RDV (black shapes), and ODBG-P-RVn (red shapes). Signal from infected cells treated with DMSO served as 100% fluorescence intensity signal for reporter assays and 100% fluorescence-positive cell counts for image-based assays. CPE inhibition was measured by determining cellular ATP levels using CellTiterGlo 2.0 assay reagent. ATP levels in uninfected cells treated with DMSO served as 100% CPE inhibition. Dose-response curves were fitted to the mean value of experiments performed in biological triplicate for each concentration in the 8-point, 3-fold dilution series using a 4-parameter non-linear logistic regression curve with variable slope. Data points and error bars indicate the mean value and standard deviation of 3 biological replicates; each colored shape/line in the legend represents an independent experiment performed in biological triplicate.

**Supplemental Figure S3.**
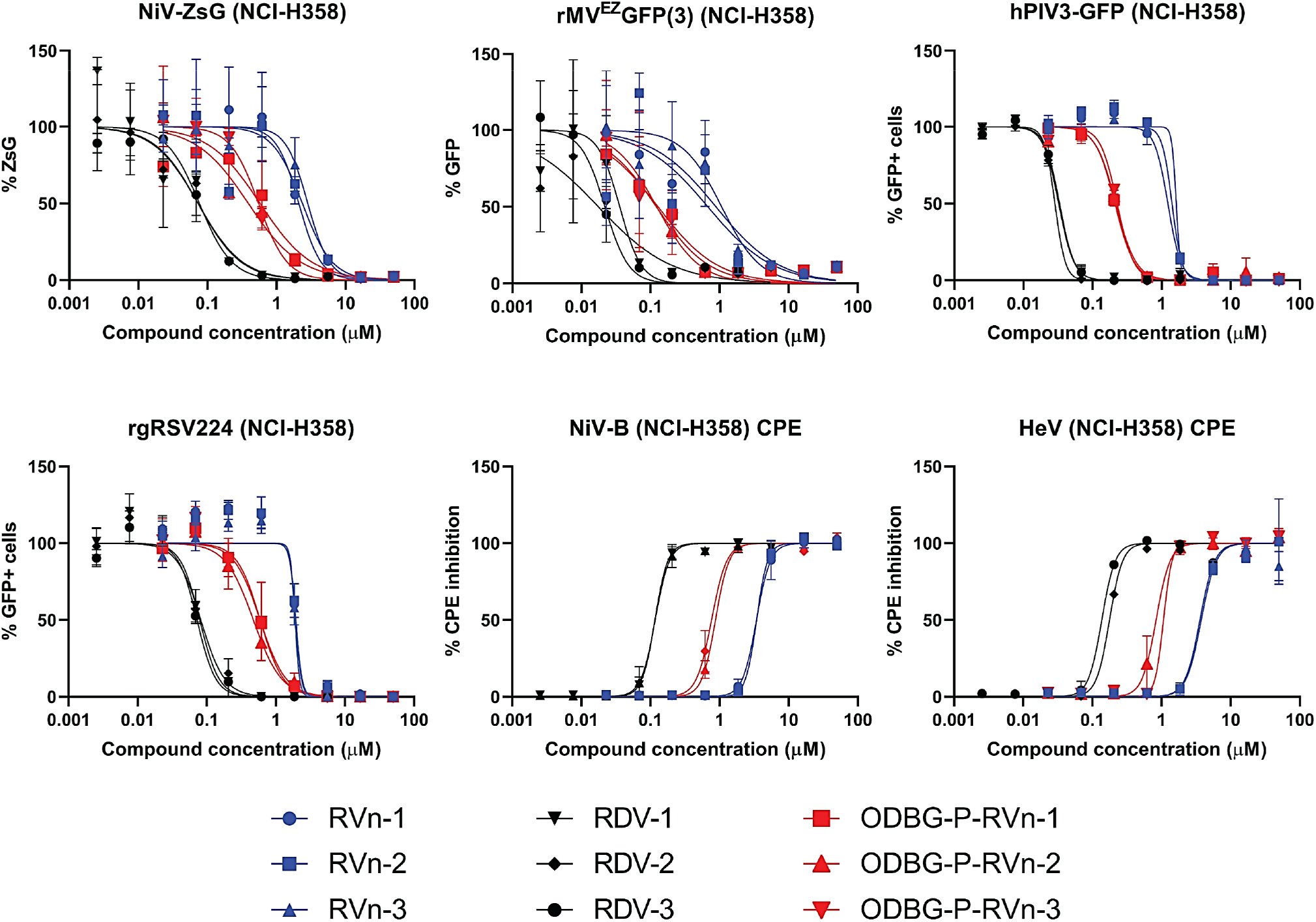
Comparison of antiviral activities of RVn, RDV, and ODBG-P-RVn in human bronchioalveolar carcinoma (NCI-H358) cells using reporter-based, image-based, and CPE assays. Representative dose-response inhibition of virus replication by RVn (blue shapes), RDV (black shapes), and ODBG-P-RVn (red shapes). Signal in infected cells treated with DMSO served as 100% fluorescence intensity signal for reporter assays and 100% fluorescence-positive cell counts for image-based assays. CPE inhibition was measured by determining cellular ATP levels using CellTiterGlo 2.0 assay reagent. ATP levels in uninfected cells treated with DMSO served as 100% CPE inhibition. Dose-response curves were fitted to the mean value of experiments performed in biological triplicate for each concentration in the 8-point, 3-fold dilution series using a 4-parameter non-linear logistic regression curve with variable slope. Data points and error bars indicate the mean value and standard deviation of 3 biological replicates; each colored shape/line in the legend represents an independent experiment performed in biological triplicate.

**Supplemental Figure S4.**
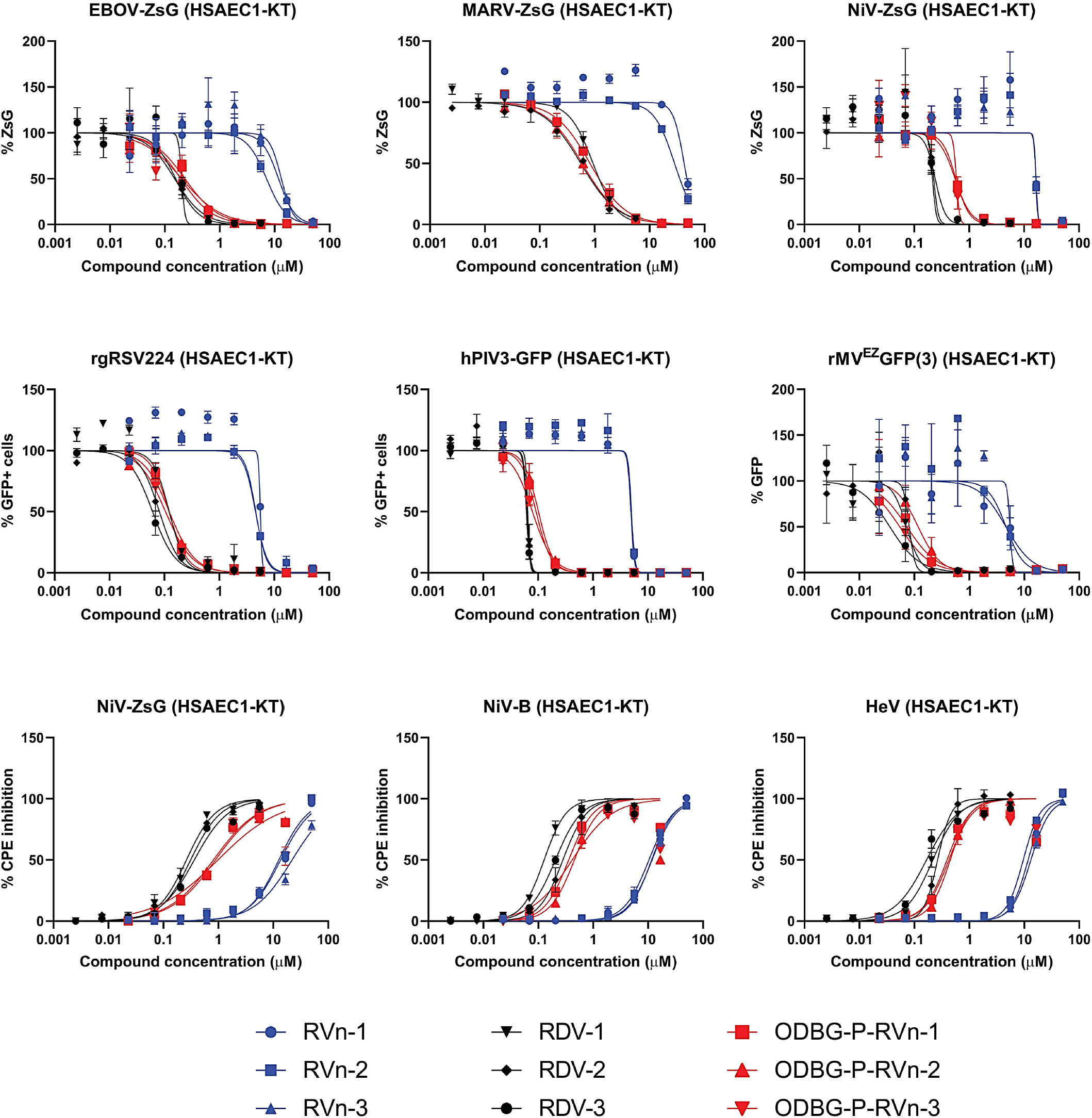
Comparison of antiviral activities of RVn, RDV, and ODBG-P-RVn in primary-like human small airway epithelial (HSAEC1-KT) cells using reporter-based, image-based, and CPE assays. Representative dose-response inhibition of virus replication by RVn (blue shapes), RDV (black shapes), and ODBG-P-RVn (red shapes). Signal in infected cells treated with DMSO served as 100% fluorescence intensity signal for reporter assays and 100% fluorescence-positive cell counts for image-based assays. CPE inhibition was measured by determining cellular ATP levels using CellTiterGlo 2.0 assay reagent. ATP levels in uninfected cells treated with DMSO served as 100% CPE inhibition. Dose-response curves were fitted to the mean value of experiments performed in biological triplicate for each concentration in the 8-point, 3-fold dilution series using a 4-parameter non-linear logistic regression curve with variable slope. Data points and error bars indicate the mean value and standard deviation of 3 biological replicates; each colored shape/line in the legend represents an independent experiment performed in biological triplicate.

